# Assessment of knowledge, attitudes and practices about antibiotic resistance among medical students in India

**DOI:** 10.1101/546713

**Authors:** Manoj Kumar Gupta, Chirag Vohra, Pankaja Raghav

## Abstract

**Background:** To reduce the magnitude of antimicrobial resistance, there is a need to change the knowledge and behavior of future prescribers regarding use and prescription of antibiotics. This can be ensured through the appropriate training of next generation doctors and medical students. But, before planning or strengthening any teaching or training program for any group, it is required to have a conclusive evidence about knowledge, attitude and practices of that group. With this background this study was conducted to assess the knowledge, attitudes and the practices of medical students in India with respect to antibiotic resistance and usage

**Methods:** It was a cross-sectional study which was done online through google forms. A semi-structured questionnaire containing a five point Likert scale was used for the data collection. The questionnaire was sent to medical students across India by sharing link through contacts of Medical Students Association of India. Respondent-driven sampling technique was also adopted for the study. Data was analyzed using SPSS v.25 and Microsoft Excel 2016.

**Results:** The overall mean score of awareness for the students was 4.36 + 0.39. As compared to first year students, mean score of awareness was significantly higher among students of all the years. A significantly better awareness was also observed among pre final year students as compared to other years. Variable practices have been observed regarding use of antibiotics among medical students.

**Conclusion:** The awareness level of medical students regarding antibiotics and its resistance was quite satisfactory. As far as attitude and practices are concerned, there is a significant need for improvements.

## Introduction

The era of antibiotics has changed the pattern of treatment and outcomes of infectious diseases. But at the same time, irrational use of antibiotics has created a havoc of antibiotic resistance.[1,2] Worldwide spread of the antibiotic resistant organisms have gradually created the threat of antimicrobial insufficiency. Patients infected with these antibiotic resistant organisms are likely to face long durations of hospital stay and require treatment with second and third line drugs, which may be more toxic and less effective.[3]

Doctors and especially future prescribers are frontline fighters against antimicrobial resistance, by rationally prescribing the antibiotics and promoting patient awareness.[4] There are sufficient evidences to support that newly licensed doctors/prescribers are not adequately trained to prescribe medications safely.[5-7] Lack of adequate training during medical degree course may be one of the reasons for that.[8]

There is a need to change the antimicrobial prescribing behavior of doctors and future prescribers to reduce the magnitude of the problem of antimicrobial resistance.[9] This can be ensured through the appropriate knowledge and training of next generation doctors and medical students.[10] Formal teaching of medical students on proper usage of antimicrobial agents may minimize the phenomena of antibiotic resistance.[11,12] But, before planning or strengthening any teaching or training program for any target group, it is required to have a conclusive evidence about baseline knowledge, attitude and practices of that group. This evidence support in devising an effective curriculum and sustainable program. With this background, this study was planned with the objective to assess the knowledge, attitudes and the practices of medical students in India with respect to antibiotic resistance and usage.

## Material and methods

It was a cross-sectional study which was done online through google forms for a period of four months from July to October 2018. Collection of primary data was done through specifically developed structured questionnaire. The questionnaire was developed based on the literature review of comparable studies.[13,14] This was validated by a pilot study on 30 medical students. To assess the awareness, students were asked to respond to 15 different statements (ten positive and five negative) on a five point Likert scale ranging from one (strongly disagree) to five (strongly agree). In this questionnaire 3 statements were related to identification of antibiotics, 3 related to role of antibiotics, 3 regarding side effects of antibiotics and 6 were related to awareness about antibiotic resistance. The questionnaire was sent to medical students across India by sharing link through contacts of Medical Students Association of India which has many students from a number of medical institutes across India as its members. They were asked to recruit further respondents into the study from their respective colleges. Thus, a respondent-driven sampling technique was also adopted for the study.

The study was approved by an institutional ethics committee. Informed consent was taken from the respondents before attempting the questionnaire. The purpose of the study was explained before starting the questionnaire and queries of all natures related to this research were invited for satisfactory explanation to ensure informed consent for participation. Data was analyzed using SPSS v.25 and Microsoft Excel 2016. Appropriate tables and graphs were prepared and inferences were drawn by applying descriptive statistics, parametric (chi-square) and non-parametric (Kruskal Wallis and Mann-Whitney U) tests.

## Results

Figure 1 depicts that a total of 474 responses were received from 103 medical colleges distributed across 22 states of India. Maximum responses were received from Rajasthan (177) followed by Delhi (59), Maharashtra (42) and Uttar Pradesh (36). (Fig. 1)

**Fig. 1:**
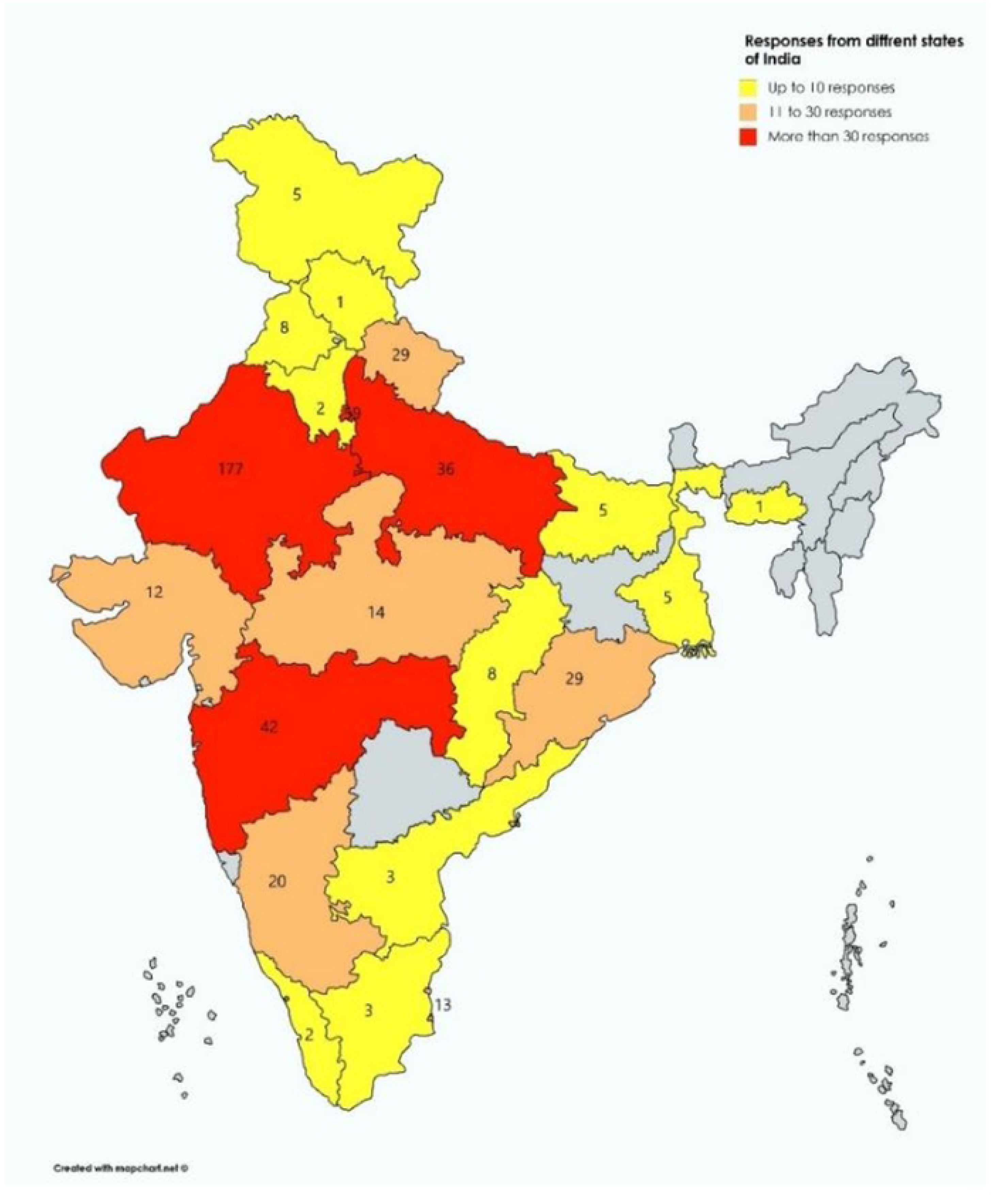
Distribution of responses from the different states of the country. As much as 32.3% of the students accepted that any of their family member was working in health related field. The descriptive of awareness about antibiotics and its resistance in terms of mean score, mean score % and median for each statement is depicted in table 1.

**Table 1:**
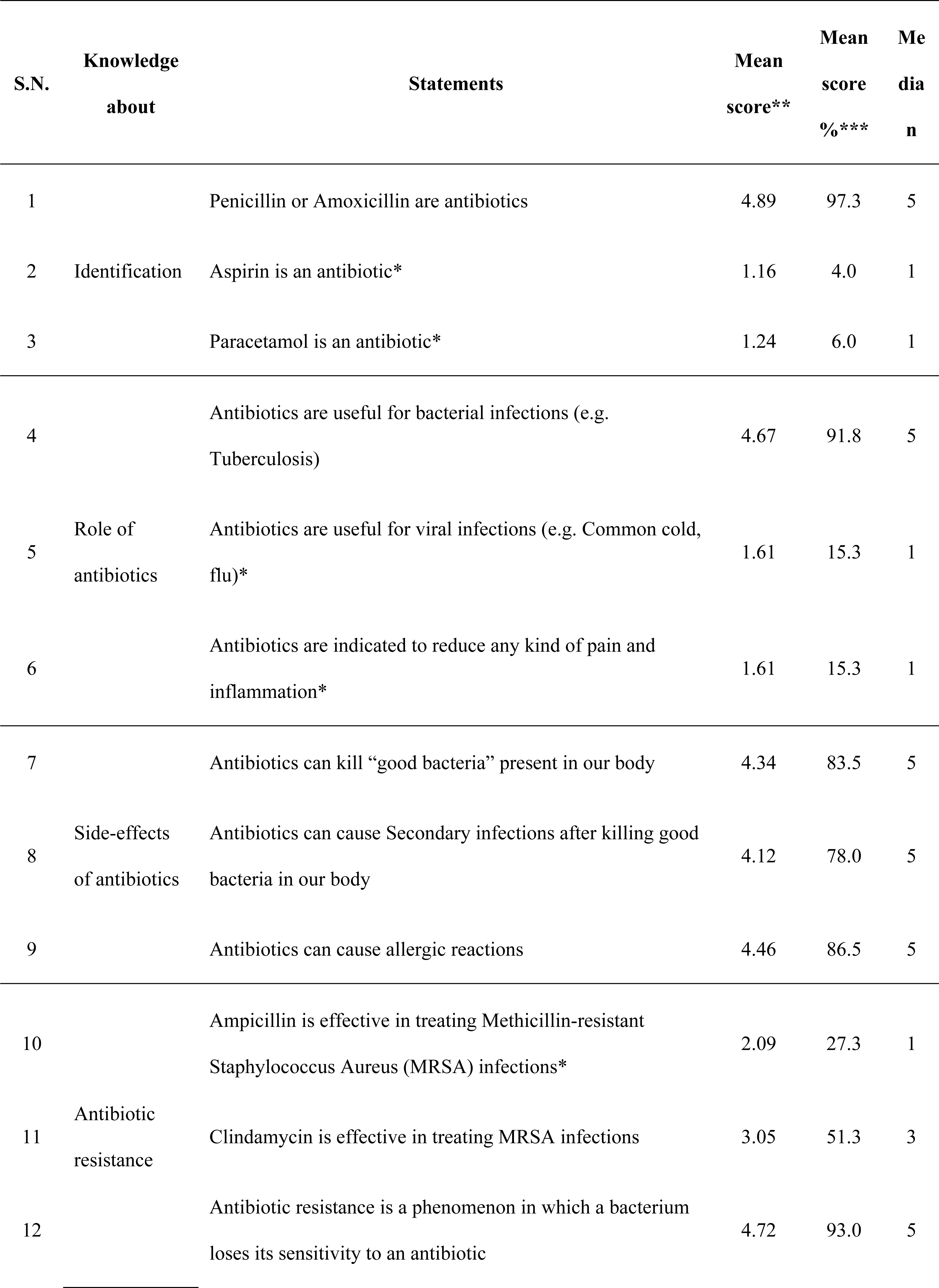
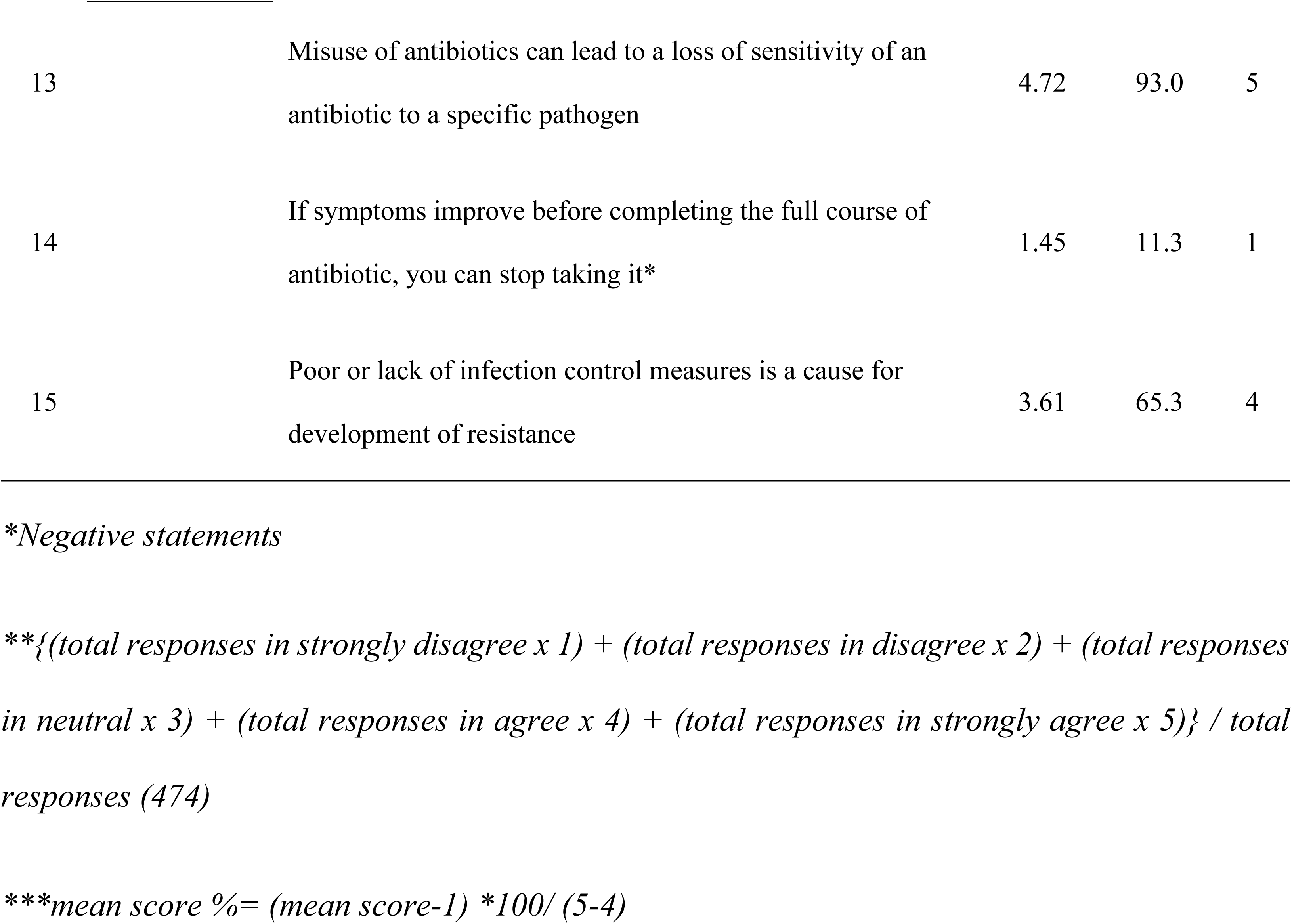
Descriptive of awareness about antibiotics and its resistance.

Table 2 depicts that maximum (40%) students were studying in pre-final year, followed by second year (27%). Reverse coding was done for five negative statements for calculating the mean score for each participant. The overall mean score was 4.36 ± 0.39. A significant difference was observed in awareness level of students according to their year of study.

**Table 2:**
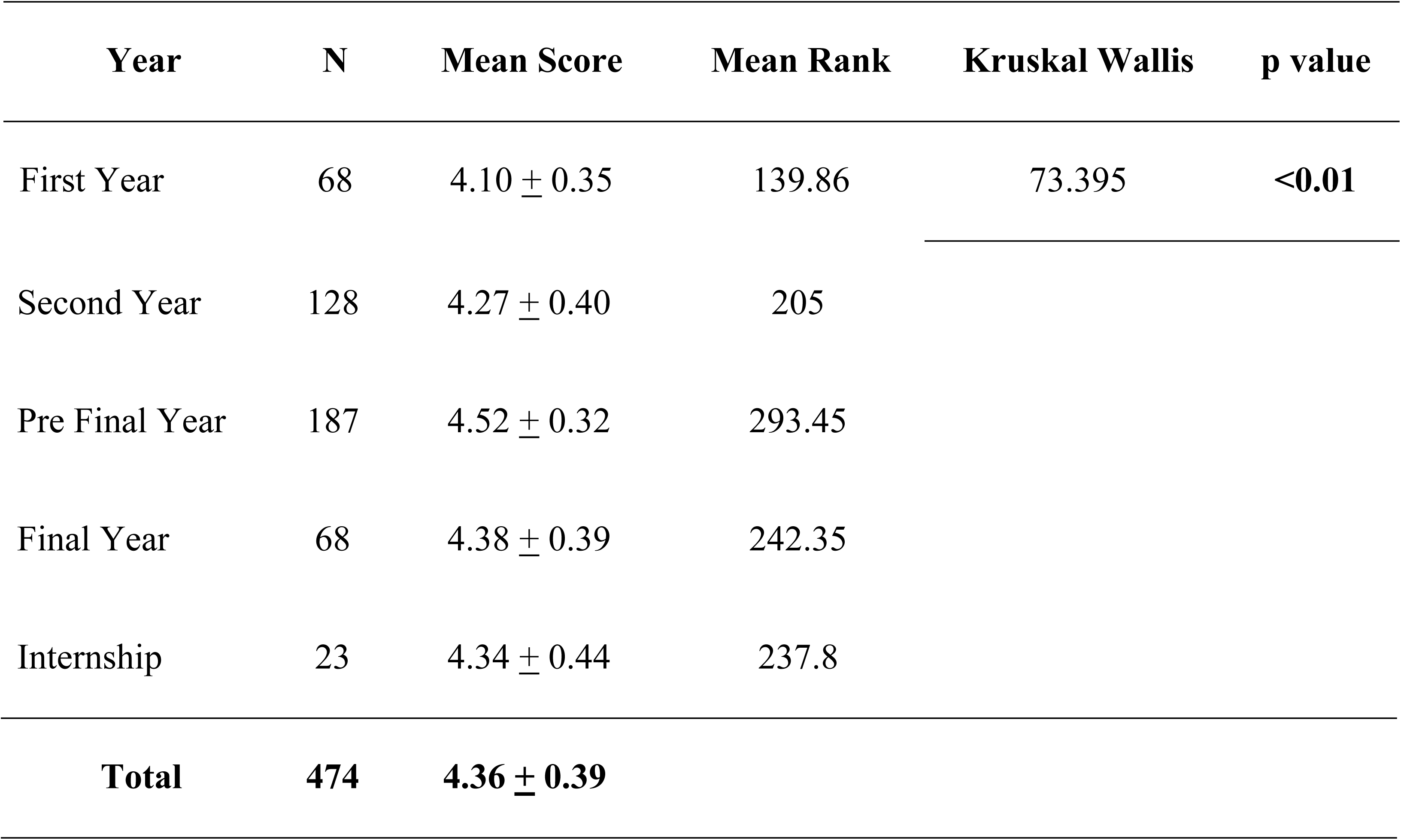
Distribution of respondents according to their year of study and mean score of knowledge.

| Year | N | Mean Score | Mean Rank | Kruskal Wallis | p value |
| --- | --- | --- | --- | --- | --- |
| First Year | 68 | $4.10 \pm 0.35$ | 139.86 | 73.395 | **<0.01** |

|  |  |  |  |
| --- | --- | --- | --- |
| Second Year | 128 | $4.27 \pm 0.40$ | 205 |
| Pre Final Year | 187 | $4.52 \pm 0.32$ | 293.45 |
| Final Year | 68 | $4.38 \pm 0.39$ | 242.35 |
| Internship | 23 | $4.34 \pm 0.44$ | 237.8 |
| **Total** | **474** | **$4.36 \pm 0.39$** |  |

To explore the exact level of difference between each category, Mann-Whitney U test was applied. (table 3).

**Table 3:**
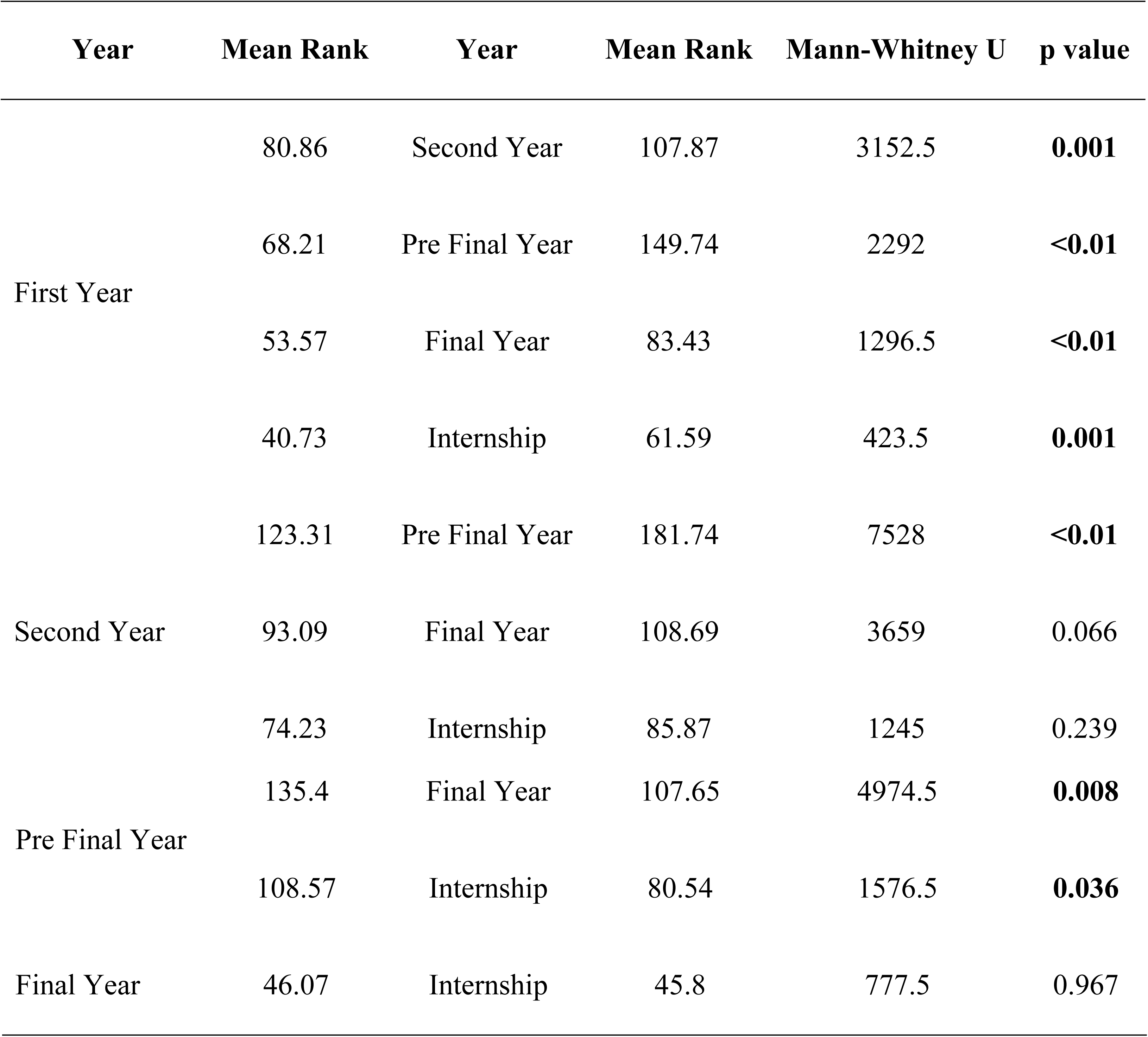
Comparison of mean score of knowledge according to their year of study.

Table 4 represents the attitude and practices of medical students regarding use of antibiotics. As much as 83.3% students have consumed antibiotics in previous year of the survey. Most of them have consumed antibiotics for once to three times in the whole year. Major source of information about antibiotics and its resistance was their degree courses.

**Table 4:** Attitude and Practices of medical students regarding use of antibiotics.

|  | Year N (%) |  |  |  |  |  |  |
| --- | --- | --- | --- | --- | --- | --- | --- |
|  | First | Second | Pre Final | Final | Interns | Total | Sig. |
| Used antibiotics in the last year | 47 (69.1) | 116 (90.6) | 153 (81.8) | 58 (85.3) | 21 (91.3) | 395 (83.3) | 0.003 |
| Number of times antibiotics used in last year (n=395) |  |  |  |  |  |  |  |
| 1 | 8 (17.0) | 28 (24.1) | 38 (24.8) | 13 (22.4) | 9 (42.9) | 96 (24.3) | 0.247 |
| 2 | 10 (21.3) | 44 (37.9) | 45 (29.4) | 15 (25.9) | 4 (19.0) | 118 (29.9) | 0.142 |
| 3 | 10 (21.3) | 20 (17.2) | 38 (24.8) | 18 (31.0) | 2 (9.5) | 88 (22.3) | 0.144 |
| 4 | 9 (19.1) | 11 (9.5) | 20 (13.1) | 9 (15.5) | 2 (9.5) | 51 (12.9) | 0.492 |
| ≥5 | 10 (21.3) | 13 (11.2) | 12 (7.8) | 3 (5.2) | 4 (19.0) | 42 (10.6) | **0.036** |
| Heard about antibiotic resistance | 68 (14.3) | 128 (27.0) | 187 (39.5) | 68 (14.3) | 23 (4.9) | 474 (100) | NA |
| Source of information about antibiotic resistance |  |  |  |  |  |  |  |
| Academic course | 27 (39.7) | 119 (93.0) | 181 (96.8) | 63 (92.6) | 21 (91.3) | 411 (86.7) | **<0.01** |
| General Practitioner | 12 (23.1) | 39 (38.2) | 36 (27.3) | 23 (40.4) | 6 (35.3) | 116 (32.2) | 0.138 |
| Internet | 24 (46.2) | 39 (38.2) | 56 (42.4) | 21 (36.8) | 9 (52.9) | 149 (41.4) | 0.665 |
| Television | 14 (26.9) | 8 (7.8) | 9 (6.8) | 8 (14.0) | 0 (0.0) | 39 (10.8) | **0.001** |
| Newspaper | 17 (32.7) | 26 (25.5) | 36 (27.3) | 13 (22.8) | 1 (5.9) | 93 (25.8) | 0.264 |
| Discussion at<br>home/school/friends | 2 (3.8) | 4 (3.9) | 3 (2.3) | 4 (7.0) | 1 (5.9) | 14 (3.9) |  |
| Usually take antibiotic for cold<br>or sore throat | 19 (27.9) | 29 (22.7) | 39 (20.9) | 21 (30.9) | 2 (8.7) | 110 (23.2) | 0.17 |
| Usually take antibiotic for fever | 30 (44.1) | 22 (17.2) | 23 (12.3) | 8 (11.8) | 2 (8.7) | 85 (17.9) | **<0.01** |
| Usually stop taking antibiotic<br>when start feeling better | 29 (42.6) | 38 (29.7) | 44 (23.5) | 17 (25.0) | 3 (13.0) | 131 (27.6) | **0.016** |
| Buy antibiotics without a<br>medical prescription | 26 (38.2) | 52 (40.6) | 85 (45.5) | 40 (58.8) | 12 (52.2) | 215 (45.4) | 0.093 |
| Keep leftovers antibiotics for<br>use in future | 45 (66.2) | 82 (64.1) | 118 (63.1) | 45 (66.2) | 15 (65.2) | 305 (64.3) | 0.987 |
| Use leftovers antibiotics without<br>consulting the doctor | 21 (30.9) | 33 (25.8) | 49 (26.2) | 27 (39.7) | 6 (26.1) | 136 (28.7) | 0.252 |
| Ever started antibiotic therapy<br>after getting consultation from<br>doctor over phone, without a<br>proper medical prescription | 30 (44.1) | 63 (49.2) | 102 (54.5) | 44 (64.7) | 13 (56.5) | 252 (53.2) | 0.14 |

## Discussion

This study has tried to explore the awareness, attitude and practices of medical students regarding antibiotic resistance. Studies advocate that medical students generally have positive attitudes about antimicrobial resistance.[15–18] There are also supportive evidence to prove that practices of self-medication is more prevalent among medical students as compared to their peer group from the nonmedical fields.[19]

### Awareness about antibiotics and its resistance

In the present study, majority of the students could correctly identify the antibiotics. Similar kinds of awareness regarding identification of antibiotics have been observed by Scaioli G. et.al. (2015) [13] among students of a school of medicine in Italy and Sharma S. et.al. (2016) [20] among medical students in Kerala, India.

The level of awareness about the fact that antibiotics are useful for bacterial infections was quite high (91.8%) among medical students. Similar level of awareness has been witnessed by many other studies.[13,14] Most of the students in the present study expressed their denial to the statement that antibiotics are useful for viral infections like common cold and flu. Comparable findings have been observed by Ahmad A et.al., (2015) among B.Sc. Pharmacy students of Trinidad and Tobago.[21] and Jorak A et.al. (2014) among medical students in Iran[22]. Many studies have reported low level of awareness among students in this regard. [4,14,23–25] A high level of disagreement (84.7%) was observed among the students to the point that antibiotics are indicated to reduce pain and inflammation in the body. Similar kind of disagreement has been observed by Sakeena MHF et.al. (2018) [25] and Jamshed SQ et.al. (2014)[11]. Contrary to these, Ajibola O et.al (2018) had observed relatively lower level of disagreement (62.8%) in this regard.[23]

The magnitude of knowledge about side effects of antibiotics was ranging from 78% to 86.5% among medical students, which is lower than the findings observed by Scaioli G. et.al. (2015) and Jamshed SQ et.al. (2014).[11,13] The knowledge about the kinds of antibiotics used for Methicillin-resistant Staphylococcus Aureus (MRSA) infections was not satisfactory among medical students. Majority of the students were aware about the mechanism of antibiotic resistance and also accepted that misuse of antibiotics can lead to antibiotic resistance to a specific pathogen. Similar findings have been reported by many studies.[13,21,25] There are other studies also in which understanding of students about the basic mechanism of antimicrobial resistance was reported quite lower.[23,26–28] As much as 88.7% disagreement was observed for the statement that, “if symptoms improve before completing the full course of antibiotic, you can stop taking it”. Mixed findings have been observed by many authors in this regard.[4,11,21,23,25,29] The mean score % of awareness for the fact that poor or lack of infection control measures is a cause for development of resistance was 63% in the present study. Similar level of awareness has been coated by other studies from different parts of the world.[26,27,30]

The overall mean score of awareness for the students was 4.36 ± 0.39. Ayepola OO et.al. (2018) found the mean knowledge score as 5.51 ± 0.14 (out of 10).[29] A significant difference was observed in awareness level of students according to their year of study. Similar kinds of observations has been made by Huang Y et.al. (2013).[14]

As compared to first year students, mean score of awareness was significantly higher among students of all the years. This is supported by the findings of Huang Y et.al. (2013).[14] A significantly better awareness was also observed among pre final year students as compared to other years. This may be due to their updated knowledge of pharmacology which they have completed in recent past year.

### Behaviors and Practices of medical students regarding use of antibiotics

Majority (83.3%) of the students had used antibiotics in previous one year of the survey. This was relatively higher than the use reported by Scaioli G et.al. (2015)[13], Sakeena MHF et.al. (2018)[25] and Ayepola OO et.al. (2018)[29].

In the present study all the students had heard about antibiotic resistance. This finding was quite higher than reported by other studies. [14,23] Major source of information about antibiotic resistance was their academic course, followed by internet, general practitioners, newspaper, television and discussions at home. Similar findings have been reported by other studies as well.[24,26]

Nearly one fourth of students gave positive affirmation regarding usually taking antibiotic for cold or sore throat. This practice was quite lower than reported by many other studies.[13,19,20,31] Huang Y et.al. (2013) observed this practice among only 13.6% of Chinese students.[14] Slightly less than one fifth students accepted that they were usually taking antibiotics for fever, which is similar to the findings reported by Ahmad A et.al. (2015). Variable practices in this regard have been reported by many other studies.[4,13,14,32]

Every one out of four students were following wrong practice of stopping antibiotics when start feeling better without completing the full course, which is similar to the findings reported by Ghaieth MF et.al (2015). Studies have reported diverse findings regarding this practice.[13,20,29,32]

Similar to other studies[19,32], in the present study slightly less than half of medical students accepted that they buy antibiotics without a medical prescription. This practice was somewhat higher than the practices reported by many other studies. [4,13,20] This self-medication practice was more prevalent among study participants of Sakeena MHF et.al. (2018)[25] and Ayepola OO et.al. (2018)[29]. Unlike other studies[13,19], no significant difference has been observed in self-medication practice among medical students as per their year of study.

Around two third of students were having practice of keeping leftover antibiotics for future use. This was similar to the findings by Ayepola OO et.al. (2018).[29] This practice was relatively lower (less than 50%) among medical students from southern part of India.[4,32]

As much as 28.7% students were using these leftovers antibiotics in future without consulting the doctor. Scaioli G et.al. (2015) found this figure as 17.7%.[13] Other studies have reported that most of the students give the leftover antibiotics to their friends, relatives or roommate without doctor’s consultation when they are sick.[20,23,32]

More than half of students accepted that they had ever started antibiotic therapy after getting consultation from doctor over phone, without a proper medical prescription. Scaioli G et.al. (2015) found this practice prevalent among around one third of students.[13] Jamshed SQ et.al. (2014) reported that 8.9% of students were having perception that prescribing antibiotics over the phone is good patient care since it can save time.[11]

## Conclusion & recommendations

The awareness level of medical students regarding antibiotics and its resistance was quite satisfactory. As far as attitude and practices are concerned, there is a significant need for improvements. Since the medical students are going to be future prescribers, it is important to have proper guidelines in medical curriculum related to use and rational prescription of antibiotics. Further there is need and scope to explore this area with large sample size and through multi-centric studies.

## Acknowledgement

We would like to acknowledge the all the medical students who participated in the study.

